# High-yield, ligation-free assembly of DNA constructs with nucleosome positioning sequence repeats for single molecule manipulation assays

**DOI:** 10.1101/2023.01.05.522917

**Authors:** Yi-Yun Lin, Tine Brouns, Pauline J. Kolbeck, Willem Vanderlinden, Jan Lipfert

## Abstract

Force and torque spectroscopy have provided unprecedented insights into the mechanical properties, conformational transitions, and dynamics of DNA and DNA-protein complexes, notably nucleosomes. Reliable single-molecule manipulation measurements require, however, specific and stable attachment chemistries to tether the molecules of interest. Here, we present a functionalization strategy for DNA that enables high-yield production of constructs for torsionally constrained and very stable attachment. The method is based on two subsequent PCR reactions: first ∼380 bp long DNA strands are generated that contain multiple labels, which are used as “megaprimers” in a second PCR reaction to generate ∼kbp long double-stranded DNA constructs with multiple labels at the respective ends. We use DBCO-based click chemistry for covalent attachment to the surface and biotin-streptavidin coupling to the bead. The resulting tethers are torsionally constrained and extremely stable under force, with an average lifetime of 60 ± 3 hours at 45 pN. The high yield of the approach enables nucleosome reconstitution by salt dialysis on the functionalized DNA and we demonstrate proof-of-concept measurements on nucleosome assembly statistics and inner turn unwrapping under force. We anticipate that our approach will facilitate a range of studies of DNA interactions and nucleoprotein complexes under forces and torques.

## INTRODUCTION

DNA is central to the storage and transmission of genetic information. Single-molecule manipulation methods have provided unprecedented insights into the mechanical properties of DNA and its processing by DNA processing enzymes [1-10]. Single-molecule force and torque spectroscopy experiments, in particular using optical or magnetic tweezers, require DNA constructs that are labeled at both ends with appropriate molecular handles to enable attachment to functionalized particles and surfaces [11, 12]. In particular, for torque and twist measurements the requirements for molecular constructs are stringent, since torsional constraint requires multiple attachment sites at both ends and completely nick-free DNA, since a single nick removes the torsional constraint by enabling free rotation.

Preparation of DNA constructs with multiple attachment sites for single-molecule torque and twist assays requires several biochemical reactions. The most frequently used protocol involves PCR, restriction reactions, and final ligation to assemble three different DNA fragments into the final construct [4, 11, 13, 14]. The central segment of the DNA molecule is unlabeled, while the two ends of the molecular construct are labeled with different moieties that enable attachment, with multiple biotin and digoxigenin (dig) labels, respectively, being a popular choice. The products of the ligation reactions are a mixture involving off-target DNA and unligated strands. Therefore, subsequent gel purification is often required to obtain the specific DNA construct. Unfortunately, these procedures lead to low yield and are prone to introduce nicks into the DNA.

To overcome the inefficiency of generating DNA through ligation based protocols, improved methods are needed to simplify the process and increase the yield. Recently, new methods have been reported to generate high-yield DNA constructs for force and torque spectroscopy experiments [15]. In particular, a ligation-free method has been reported that achieves high yield of torsionally constrained DNA and efficiently incorporates different labels at the end of DNA [16]. The strategy is based on two PCR-synthesized DNA constructs that are used as “megaprimers” with either biotin or digoxigenin labels to amplify the target DNA in a final PCR step. The megaprimer-based PCR reaction can create torsionally constrained DNA without ligation and restriction reactions. However, an additional requirement for biomechanical characterization at the single-molecule level is efficient and stable tethering of the biomolecules of interest. The commonly used attachment strategy relying on the binding of biotin to streptavidin-coated beads and the binding of digoxygenin (dig) labeled nucleotides to a surface coated with antibodies against digoxygenin (anti-dig) is rapid and reliable. The biotin-streptavidin interaction exhibits good force stability, despite being non-covalent, and can be optimized through engineered streptavidin variants [17-19]. Unfortunately, the anti-dig/dig interaction has much lower stability under forces > 10 pN [20, 21]. Going beyond non-covalent attachment, introduction of one or more covalent linkages for biomolecular attachment provides force stabilities up to nN [22]. Several approaches for covalent attachment have been developed [22-25] and provide stability for long measurements and experiments over a broad force spectrum [8, 26-30]. For DNA attachment via dibenzocyclooctyne (DBCO), covalent binding to an azide-functionalized surface has been developed. This copper-free click chemistry method is specific, highly efficient and yields tethers that are able to withstand very high forces (>100 pN) [25].

Here, we present a novel attachment protocol that combines the high-yield enabled by the megaprimer approach with the supreme force stability afforded by covalent DBCO/azide coupling for anchoring DNA in tweezers experiments. Our strategy is based on DNA constructs with multiple biotin and DBCO labels at each end, respectively, and assembly via ligation-free PCR. The constructs enable torsionally constrained coupling and excellent force stability. The high yield of the approach enables us to obtain sufficient material for downstream biochemical preparation, in particular for nucleosome reconstitution.

The eukaryotic genome is highly compacted within the nucleus to form chromatin. Nucleosomes are the basic units of chromatin. Canonical nucleosome core particles are composed of two copies of H2A, H2B, H3 and H4 assembled into a histone octamer that is wrapped by 147 bp of DNA [31, 32]. In order to study nucleosome interactions *in vitro*, nucleosome-positioning sequences with a high affinity for histone octamers are often used and nucleosomes can be assembled by reconstitution, typically via salt dialysis. In particular, the “Widom 601” sequence [33] has become widely used for studies of nucleosome structure and function *in vitro*.

To reconstitute nucleosomes *in vitro*, at least 1 μg or 10 ng/μL DNA with appropriate positioning sequences [34-36] are required, to enable efficient handling and high enough concentrations for the assembly reaction. PCR amplifications of repetitive DNA and with megaprimers that contain different chemical labels are problematic as they can generate undesired products and low yield. Therefore, we optimized reaction conditions and tested different polymerases to obtain high yields. Finally, we characterized the mechanical properties of our new DNA constructs and the products of the nucleosome reconstitution. For both DNA and nucleosome constructs, we achieved reliable tethering and found mechanical signatures in good agreement with previous studies.

## MATERIALS AND METHODS

### DNA preparation

We used the plasmid pFMP218 [37] as template to produce DNA constructs with nucleosome positing sequences. pFMP218 is a custom-built plasmid (provided by Prof. Felix Müller-Planitz, TU Dresden, Germany) from a pUC18 backbone and with 3 repeats of Widom 601 sequences inserted. The final design yields a 2823 bp DNA molecule with a 2055 bp central unlabeled segment, flanked by two ∼380 bp labeled regions. We generated the 2823 bp DNA construct from pFMP218 as linear template for subsequent assembly by PCR with Phusion Hot Start polymerase (follow the vendor’s protocol) using forward primer 5’-GCAGAAGTGGTCCTGCAACT-3’ and reverse primer 5’-CCGGATCAAGAGCTACCAAC-3’.

Functionalized handles (the “megaprimers”; Figure 1a) were obtained by PCR amplification. We tested both Taq polymerase (New England Biolabs (NEB), Ipswich, MA USA) and KOD Hot Start polymerase (Novagen, Darmstadt, Germany) to prepare the functionalized handles and found both to give good yields of functionalized DNA product (Supplemenatry Figure S1). For both polymerases, we ran PCR reactions with added biotin-16-dUTP or 5-DBCO-(PEG)_4_-dUTP (Jena Biosciences GmbH, Jena, Germany), respectively. For Taq polymerase, the PCR reactions used 0.3 μM forward primer 5’-GCAGAAGTGGTCCTGCAACT-3’, 0.3 μM reverse primer 5’-CGCCGCATACACTATTCTCA-3’, 6 ng template DNA, and different amounts of biotin-16-dUTP (0.16, 0.08, or 0.04 mM final concentration) in 20 μL 1 x Taq Master Mix (which contains a final concentration of 0.2 mM of each unmodified dNTP). The other PCR solution contains 0.3 μM forward primer 5’-CCGCTTACCGGATACCTGTC-3’, 0.3 μM reverse primer 5’-CCGGATCAAGAGCTACCAAC-3’, 6 ng template DNA, and different amounts of 5-DBCO-(PEG)_4_-dUTP (0.16, 0.08, or 0.04 mM final concentration) in 20 μL 1 x Taq Master Mix.

**Figure 1.**
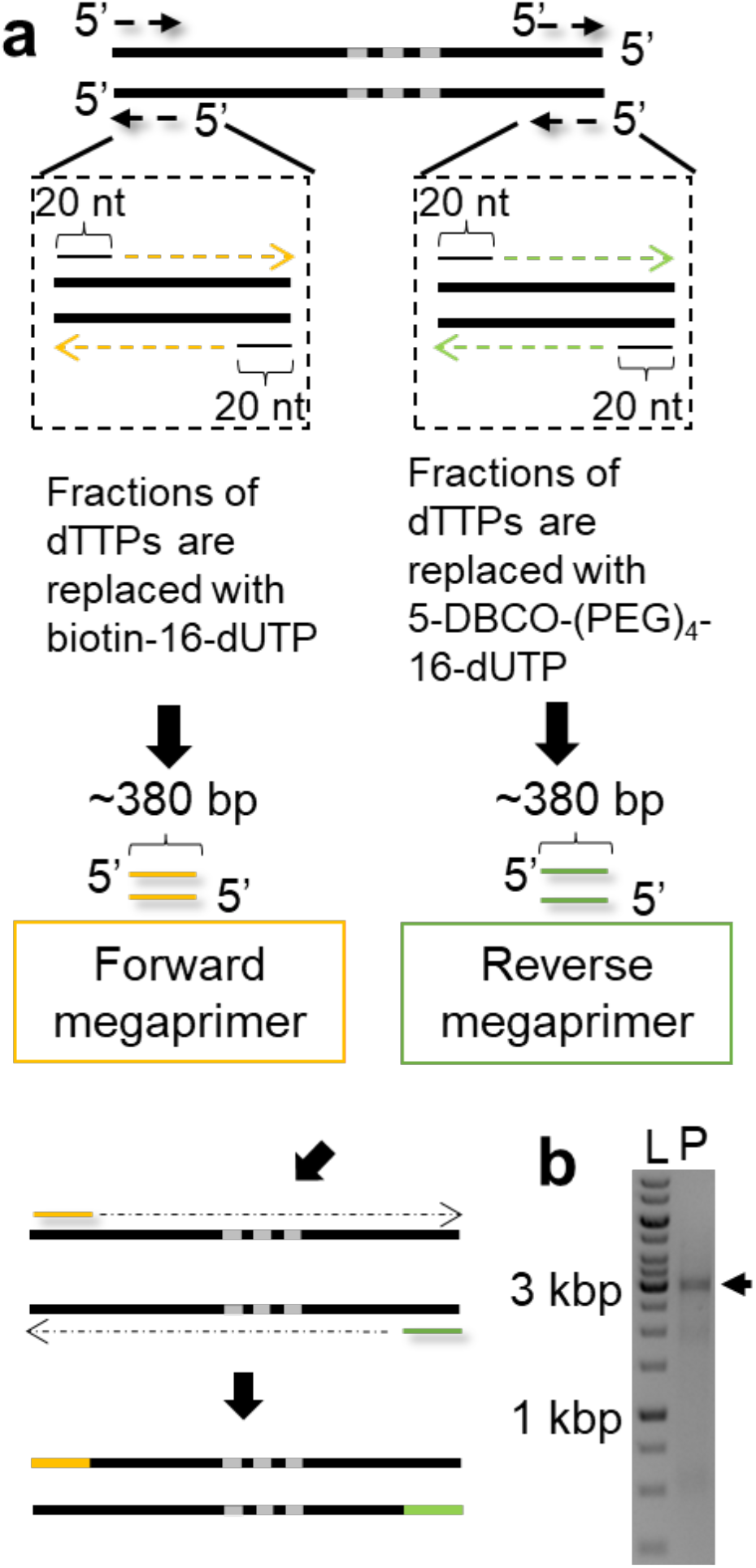
Ligation-free megaprimers PCR-based DNA assembly method. (a) Schematic of the ligation-free method to synthesize torsionally constrained DNA. Briefly, two sets of 20-bp ssDNA primers and linearized templates are used in PCR reactions to make two multi-labeled ∼380 bp DNAs that become the megaprimer. The two megaprimers are labeled with biotin and DBCO, respectively. A linearized template with three repeats of the Widom 601 sequence (shown schematically as grey bars) is used for subsequent PCR to get the final 2823 bp PCR construct. (b) Visualization of the PCR result by gel electrophoresis. The left lane (“L”) has a DNA size ladder (1 kb+, NE Biolabs). The black arrow in the left lane (“P”) indicates the position of PCR product.

For KOD Hot Start polymerase, we followed a previous protocol [16]. We used the same DNA template and primers as for Taq polymerase and incorporated labeled dUTPs by replacing dTTPs in the PCR reaction mix [38] using either 25% or 50% Biotin-16-dUTP or DBCO-(PEG)_4_-dUTP (the final concentrations are 0.05/0.15 and 0.1/0.1 mM modified dUTP/unmodified dTTP), respectively.

The functionalized PCR products are subsequently used as megaprimers to amplify the desired DNA substrate. We used KOD Hot Start polymerase for this PCR reaction. Instead of following the vendor’s protocol, we changed the reaction conditions to optimize the purity and yield of our target DNA (Supplementary Figure S2). Our final protocol uses KOD Hot Start polymerase with a reaction solution containing 1.5 mM MgCl_2_, 0.16 mM x 4 dNTP, 1x KOD Hot start buffer, 7.5% DMSO (New England Biolabs), 200 ng forward megaprimer, 200 ng reverse megaprimer, 50 ng linear template from pFMP218, and 0.5 μL (1 unit/μL) KOD Hot start polymerase in 100 μL reaction volume. We used the following PCR cycling parameters: Initial denaturation at 95 ºC for 2 min; 35 cycles of denaturation at 95 ºC for 20 s, annealing at 60 ºC for 10 s, and elongation at 70 ºC for 65 s. The final cycle was followed by extension at 70ºC for 1 min. The PCR products were purified by using the QIAquick PCR Purification Kit (Qiagen, Hilden, Germany) after each step of PCR amplification.

We also tested PCR reactions using a 5x smaller reaction volume and 5x higher final concentrations of megaprimers and template DNA, using 10% DMSO and 1x KOD Hot start polymerase MasterMix (Novagen, Darmstadt, Germany) (Supplementary Figure S3). The smaller final volume facilitates downstream processing and the higher megaprimer concentration has been suggested to increase yield [39, 40]. While the PCR reactions at higher concentrations yield functionalized DNA constructs, we find more off-target products (Supplementary Figure S3) and greater variability in yield. Therefore, we recommend the reaction described above with a final reaction volume of 100 μl.

### Nucleosome reconstitution

Nucleosomes were assembled on the labeled DNA construct assembled using the megaprimer protocol outlined in the previous section. Recombinant human histone octamers were purchased from EpiCypher (Durham, North Carolina). Samples were prepared via salt gradient dialysis following the protocol described previously [41]. In brief, 2.7 μg DNA (∼150 nM of 601 motifs) and 0.6 μg histone octamer (170 nM) in 30 μL high-salt buffer (10 mM Tris-HCl, pH 7.6, 1 mM EDTA, 0.1 % (w/v) Triton-X100, and 2 M NaCl) were incubated in a dialysis chamber (Slide-A-Lyzer MINI Dialysis Devices, 3.5K MWCO, Thermo Scientific) at 4°C. Then, the dialysis chamber was transferred to a glass beaker with 300 mL high-salt buffer and 300 μL β-mercaptoethanol. 3 L low-salt buffer (10 mM Tris-HCl, pH 7.6, 1 mM EDTA, 0.1 % (w/v) Triton-X100, and 50 mM NaCl) and 300 μL β-mercaptoethanol were transferred to the high-salt buffer overnight at 4°C (at least 16 h). The buffer exchange is achieved with a peristaltic pump to slowly introduce the low-salt buffer into the beaker with the dialysis chamber. Finally, we placed the dialysis chamber into 1 L low-salt buffer with 300 μL β-mercaptoethanol for 1-2 h.

### Magnetic tweezers setup

We used a custom-built MT setup described previously [42]. The setup employs a pair of 5 × 5 × 5 mm^3^ permanent magnets (W-05-N50-G, Supermagnete, Switzerland) with a 1 mm gap in vertical configuration [43]. We used a DC-motor (M-126.PD2, P1, Germany) to control the distance between magnets and the flow cell. A LED (69647, Lumitronix LED Technik GmbH, Germany) was used for illumination. We used a 40x oil-immersion objective (UPLFLN 40x, Olympus, Japan) and a CMOS sensor camera with 4096 × 3072 pixels (12M Falcon2, Teledyne Daisa, Canada) to image a field of view of 400 × 300 μm^2^. Images were recorded at 58 Hz and transferred to a frame grabber (PCIe 1433; National Instruments, Austin TX). Images are tracked in real-time with custom-written tracking software (Labview, National Instruments) to extract the (x,y,z) coordinates of all beads. The objective is mounted on a piezo stage (Pifoc P726. 1CD, PI Physikinstrumente) to build a look-up table (LUT) for tracking the bead z-position. With a step size of 100 nm, the LUT was generated over a range of 10 μm. Set up control and bead tracking used Labview routines described previously [44].

### Flow cell assembly and preparation

Flow cells were assembled from two microscope cover slips with a parafilm spacer. The bottom coverslip (24 × 60 mm, Carl Roth, Germany) was treated with 2 % APTES to generate an aminosilanized surface. Before flow cell assembly, 5000x diluted stock solution of polystyrene beads (Polysciences, USA) in ethanol (Carl Roth, Germany) was deposited on the amino-coated coverslip and then slowly dried. These immobile surface-bound beads serve as reference beads for drift correction. The bottom coverslip was aligned with a pre-cut parafilm and a top coverslip with two small holes for inlet and outlet. Then the assembled flow cell was baked at 80 °C for 1 min.

### DNA or polynucleosome anchoring for magnetic tweezers experiments

Following flow cell assembly, 50 mM each of Azide-(PEG)_4_-NHS (Jena Biosciences GmbH, Jena, Germany) and Methyl-(PEG)_4_-NHS (Life technologies) in 1 x PBS were introduced and incubated for 1 h [25]. To prepare the solution, 100 mg Azide-(PEG)_4_-NHS or Methyl-(PEG)_4_-NHS were dissolved in 100 μL DMSO, respectively, to prevent hydrolysis of the NHS ester during storage at –20°C. We added 1 x PBS buffer to adjust to a final concentration of 100 mM for both Azide-(PEG)_4_-NHS and Methyl-(PEG)_4_-NHS, respectively, and then mix equal volumes. The mixture is quickly filled into the incubation chamber for surface passivation to avoid hydrolysis. We found that the addition of salt in HEPES buffer (10 mM HEPES, pH 7.6) is necessary to immobilize DNA or polynucleosomes on the surface of the flow cell. Therefore, we mixed our DNA or polynucleosome sample in measurement buffer MB1 (MB1; 10 mM HEPES pH 7.6, 100 mM KCl, 2 mM MgCl_2_, 0.1 % Tween-20).

Next, the flow cell was flushed with 500 μl MB1. DNA or polynucleosomes were dissolved in 100 μl MB1, flushed into the flow cell and incubated for 1 h. Afterwards, we rinse with MB2 buffer, which consists of MB1 supplemented with 0.1% (w/v) bovine serum albumin (Carl Roth, Germany). The flow cell was rinsed with MB2 to flush out unbound DNA or nucleosomes. Subsequently, we flow in 1% casein for nucleosome samples or 1.5% (w/v) bovine serum albumin for DNA sample in MB2 into the flow cell, incubate for 1 h to minimize nonspecific interactions, and then flush with MB2. Finally, we flush in streptavidin-coated M270 beads (Dynabeads, Invitrogen) and incubate in the flow cell for 10 min. Subsequently, unbound beads are flushed out with 2 ml MB2.

### AFM sample preparation, imaging, and analysis

We follow the previously published protocol to prepare samples for AFM imaging [45-48]. Briefly, reconstituted nucleosomes were incubated in 200 mM NaCl and 10 mM Tris-HCl, pH 7.6, for 1 min on ice and then deposited on poly-L-lysine (0.01% w/v) coated muscovite mica for 30 s, followed by 20 ml Milli-Q water rinsing and drying with a gentle stream of filtered N_2_ gas. AFM imaging was performed on a Nanowizard Ultraspeed 2 (JPK, Berlin,Germany) with AFM cantilevers, FASTSCAN-A (resonance frequency 1400 kHz, spring constant 18 N/m; Bruker), for high-speed imaging in air. All AFM images were acquired in tapping mode at room temperature. The scans were recorded at 3 Hz line frequency over a field of view of 3 μm x 3 μm at 2048 × 2048 pixels. For image processing, Scanning Probe Image Processor (SPIP v6.4; Image Metrology) was employed. Image processing involved background correction by using global fitting with a third-order polynomial and line-by-line correction through the histogram alignment routine.

### Model for the step size distributions

When subjecting the megaprimer-DNA tethers to constant forces, we observe step-wise increases in the tether length prior to tether rupture. Here we describe a simple, minimal model to account for the experimentally observed step size distribution. Since the covalent linkages used in our protocol are expected to be very force stable, we assume that only the biotin-streptavidin bonds dissociate under the constant loads exerted in the magnetic tweezers assay and cause the observed steps on the time scale of our experiments [19, 49]. As a starting point, we used the DNA sequence for the biotin megaprimer assembly. In our reaction, a fraction of the thymidine moieties in the DNA sequence is replaced by biotinylated uridine. However, it is important to note that not all thymidine moieties in the sequence will form bonds to the magnetic bead since i) the PCR reaction contains a mix of labeled and unlabeled nucleotides, ii) the polymerase might incorporate labeled and unlabeled nucleotides with different probabilities [50, 51], iii) not all labeled nucleotides will bind to a streptavidin on the bead due to steric constraints, amongst other reasons. We assume that the incorporation and subsequent binding to streptavidin of functionalized nucleotides are random and occurs with probability *P*_label_ that we treat as a fitting parameter. We used a Monte Carlo approach whereby PCR reactions are simulated by randomly incorporating labels at T positions in the sequence with probability *P*_label_. To convert the simulated label positions in the sequence to step sizes, we further assumed that streptavidin can only bind labels that are at least 10 bp apart, corresponding to a minimal physical distance of ∼3 nm, since it is unlikely that two streptavidin tetramers [52] would bind within one helical turn. Simulated step size distributions are obtained by converting the distances between subsequent incorporated and streptavidin bound labels from bp to nm using a conversion factor of 0.34 nm/bp, corresponding to the crystallographic length of double-stranded DNA. The experimentally determined step size distribution is compared to the experimental data using an unweighted χ^2^-criterion. In addition, we compare the mean step sizes of the simulated and experimental data.

## RESULT

Our protocol for the construction of end-labeled DNA constructs for single-molecule experiments has two key PCR steps (Figure 1a). In the first step, regular primers are used in PCR reactions that include modified nucleotides to generate ∼380 bp labeled DNA constructs. These labeled DNA constructs are used as “megaprimers” in a second PCR reaction with regular nucleotides to generate the final ∼kbp DNA constructs with labeled ends.

### Generation of labeled megaprimer constructs by PCR

We assembled 380 bp dsDNA labeled megaprimer constructs by PCR. In the PCR reaction to assemble functionalized megaprimers, we used 20 nt forward and reverse primer, pFMP218 as template, and a non-proofreading Taq polymerase or KOD Hot Start polymerase (see Methods). We added Biotin-16-dUTP or DBCO-(PEG)_4-_dUTP to the PCR reaction, up to 50 % of modified dUTP. Gel analysis shows that the PCR reactions with labeled nucleotides yield single bands (Supplementary Figure S1). As the percentage of modified dNTPs increases, a decrease in amplicon mobility is observed. This is consistent with the previously observed change in mobility due to the bulky side chains introduced by biotin-dUTP derivatives [53]. We similarly attributed the mobility shift of the DBCO-(PEG)_4-_dUTP substitution to the bulky side chain. We find that amongst the tested conditions, the mobility of the amplicons is lowest (and thus amount of incorporated labels highest) for KOD Hot Start polymerase with 50% dTTP substitution (Supplementary Figure S1). We, therefore, use the megaprimers generated using the combination of KOD polymerase and 50% dTTP substitution in the subsequent steps.

### Megaprimer PCR reaction to generate labeled DNA constructs for single-molecule measurements

The megaprimer approach has originally been used for site-direct mutagenesis. More recently, the approach has been expanded by using biotin-labeled and dig-labeled megaprimers to generate DNA constructs with multiple labels at both ends by PCR amplification [16]. We optimized conditions to assemble labeled DNA construct with arrays of nucleosome positioning sequences by PCR. A biotin-labeled megaprimer is used in this second PCR step as the forward primer and a DBCO-labeled megaprimer is used as reverse primer, to generate final DNA constructs with functionalized ends (Figure 1). To suppress off-target products, we prepared a linear template by PCR amplification. As the template, we used a 2823 bp plasmid DNA comprising three Widom 601 sequences. We followed the megaprimer method described previously [16] as a starting point, which uses KOD Hot Start polymerase, which is thermostable and designed to amplify difficult amplicons. We optimized PCR reaction conditions by adding DMSO and decreasing the polymerase concentration (Supplementary Figure S2 and S3 and Methods). To test the applicability of the customized protocol, we combined different substitutions of megaprimers to process PCR reactions. In our final protocol, we use megaprimers prepared with 50% biotin-dUTP and 50% DBCO-dUTP substitution (Supplementary Figure S2).

To test the reproducibility of our protocol, we performed 5 repeats of the PCR amplification for different megaprimer combinations by following our optimized protocol. We find that each repeat of the experiments successfully amplified the target products (Supplementary Figure S2). To assess the purity from the PCR amplifications, we quantified the target product intensity relative to the integrated lane intensity (Supplementary Figure S2). The average purities from different combinations of megaprimer show similar values for different megaprimers used and are >50% in all cases tested. In summary, our optimized megaprimer PCR approach enables reproducible amplification of high purity DNA constructs with specific modifications at both ends. The yield and purity are considerably higher than what is achieved in previous, ligation-based protocols, which is critical in particular for nucleosome assembly on the functionalized DNA. In addition to the DNA constructs with three Widom 601 nucleosome positioning sequences, we also generated a functionalized DNA construct with one Widom 601 (Supplementary Figure S4) and a longer 6.6 kbp DNA without nucleosome positioning sequences (Supplementary Figure S5), using the megaprimer approach with biotin and DBCO labels. In both cases we obtain torsionally constrained DNA tethers suitable for magnetic tweezers experiments (Supplementary Figure S4 and S5), demonstrating the versatility of the approach.

### Force response of megaprimer DNA constructs in magnetic tweezers

To obtain a high density of labels on the DNA, we use 50% Biotin-dUTP and 50% DBCO-dUTP megaprimers for amplification and test the resulting labeled DNA construct in our single-molecule magnetic tweezers set up (Figure 2a and Methods). In the magnetic tweezers flow cell, the 2823 bp DNA tethers are anchored with copper-free click chemistry [25] to the surface and via biotin-streptavidin coupling to magnetic beads. We track the position of the magnetic beads while applying calibrated forces [43, 54]. From the mean extension as a function of applied force, we obtain force-extension curves (Figure 2b). As expected, the force-extension behavior of DNA closely follows the inextensible worm-like chain (WLC) model until 10 pN. At forces > 60 pN, the force-extension response of the DNA tethers exhibits two different behaviors: some molecules still behave similar to the worm-like chain model, while others exhibit overstretching behavior at about 65 pN, which corresponds to the characteristic signature of stretching torsionally unconstrained double-stranded DNA [2, 55, 56]. The absence of an overstretching transition in our force range for some DNA tethers is consistent with the expectation for fully torsionally constrained DNA, for which the overstretching transition is suppressed at forces below 110 pN [2, 56].

**Figure 2.**
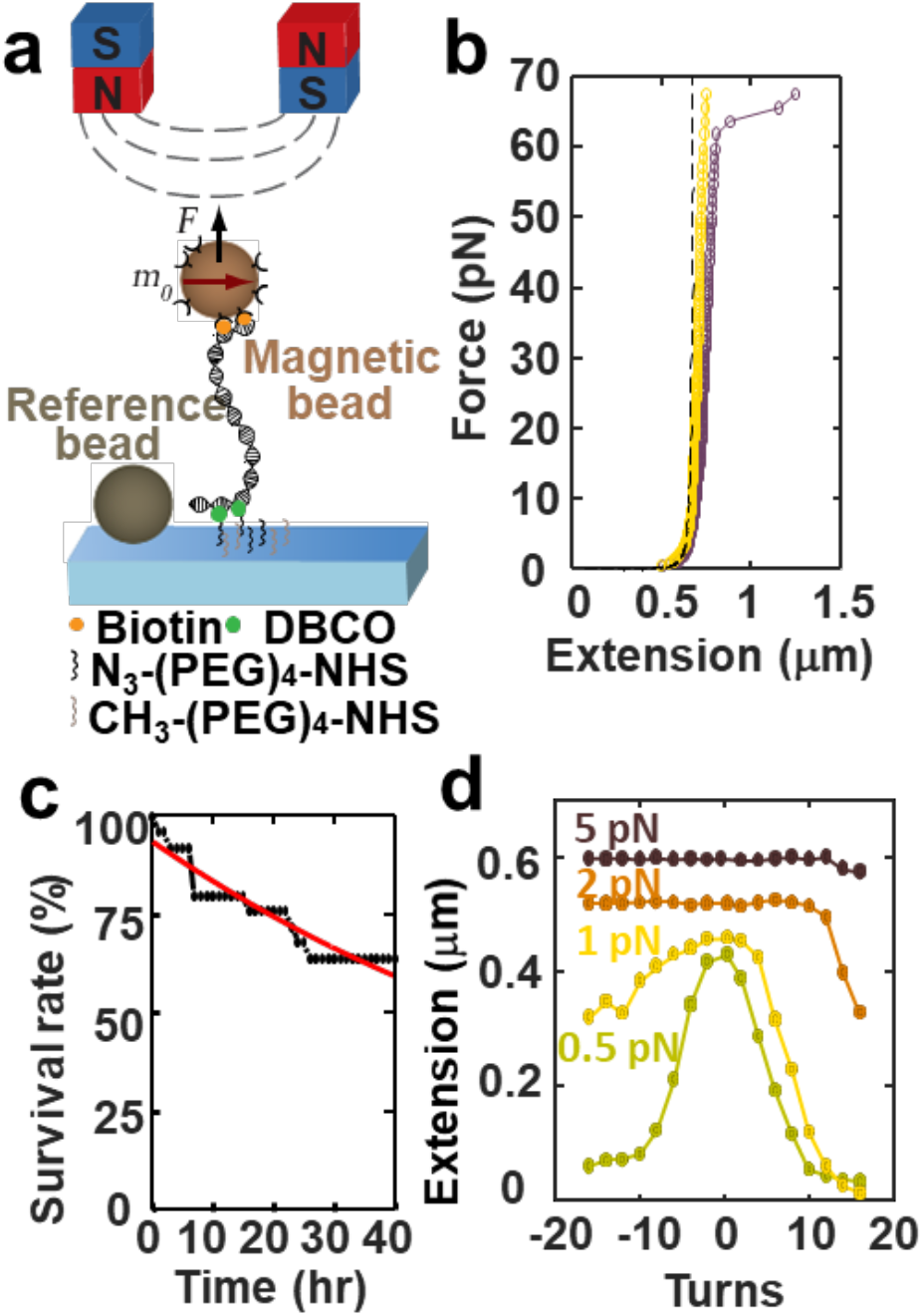
Force spectroscopy experiments on megaprimer generated DNA constructs in magnetic tweezers. (a) Schematic of the magnetic tweezers set-up. The flow cell surface is functionalized with azide-(PEG)_4_-NHS. The DNA construct has two handles, one labeled with multiple biotins and the other with multiple DBCOs. The biotins bind to multiple streptavidins that coat the magnetic bead and DBCOs at the other end form the covalent bond with the azide group. (b) Force-extension curve of DNA molecules anchored as shown in panel a. Representative force-extension measurements of torsionally constrained (yellow), unconstrained DNA molecules (brown), and co-plot of the inextensible WLC model with a bending persistence length of 45 nm (black dashed line). Torsionally unconstrained DNA exhibits the overstretching transition near 65 pN; the torsionally constrained molecule does not overstretch at 65 pN. (c) Force-stability experiment: DNA tethers are subjected to a constant force of 45 pN and tether ruptures recorded. The red line is an exponential fit to the data with a mean lifetime of 60 ± 3 h. (d) Extension-rotation curves at constant forces of 0.5, 1, 2, 5 pN indicate that the DNA construct is torsionally constrained and shows the well-known response of double-stranded DNA [4, 5].

To determine the lifetime of our DNA tethers under mechanical load, we subjected them to a constant force of 45 pN and recorded the position traces of the beads until rupture of the molecular tethers. We analyzed the lifetimes until rupture and found an exponential lifetime distribution with a fitted mean lifetime of *τ*_1_ = 60 ± 3 h (Figure 2c). Our DNA construct is modified with multiple biotin and DBCO labels at the opposite ends. The DBCO based coupling has been reported to provide force stability up to nN [22, 25]. In contrast, while the streptavidin-biotin bond has an extraordinarily high affinity (*K*_*d*_ ∼ 10^−14^ M), the interaction is non-covalent and can be broken under external forces well below 1 nN [17, 19, 57]. The tetrameric structure of streptavidin and its non-specific coupling for commercially available beads means that different force-loading geometries are possible, which gives rise to a broad range of multi-exponential lifetimes [17-19]. Previous work has shown that an engineered monovalent variant of streptavidin in the most stable geometry (1SA) exhibits a lifetime *τ*_1_ = 11.2 ± 0.4 h at 45 pN for a single biotin-streptavidin bond. We attribute the fact that we observe an even longer lifetime than what was found for the single, engineered streptavidin at the same force, despite using commercially available beads without the optimized streptavidin, to the fact that our tethers feature multiple biotin labels that can bind to the bead and, therefore, significantly increase the overall lifetime under force. Similarly, Janissen *et al*. have investigated different DNA constructs by using magnetic tweezers [21]. They found that the lifetime under a force of 45 pN for traditional biotin and digoxigenin-based DNA anchoring is only ∼7 min. If the digoxigenin is replaced by covalent anchoring while retaining the single biotin-streptavidin linkage at the other end, the lifetime increases to ∼3 h, highlighting the dramatic increase in force stability afforded by replacing the dig-antidig coupling to the surface with a covalent attachment approach. The fact that we achieve an approximately 20-fold longer lifetime compared to Janissen *et al*. might be due to their lower label density or the fact that they use neutravidin instead of streptavidin [13, 21].

### Torsional response of megaprimer DNA constructs in magnetic tweezers

To determine whether our attachment protocol with multiple labels at both ends enables supercoiling experiments on DNA tethers, we systematically under- and over-wound the DNA by rotating the magnets from –16 to 16 turns at forces of 0.5, 1, 2, and 5 pN (Figure 2d). Below 1 pN, the rotation curve of DNA behaves symmetrically and the extension of DNA decreases on overwinding or underwinding past the buckling point [4, 5, 13]. At 1 and 2 pN, the extension-rotation response of the DNA is asymmetric, due to torque-induced melting of the DNA [58]. Overall, our measurements recapitulate the well-known extension-rotation response of double-stranded DNA. We typically obtain 50 % of supercoilable (i.e. fully torsionally constrained) DNA tethers. For the 6.6 kbp DNA construct without nucleosome positioning sequences, we performed additional experiments using high-speed tracking at 1 kHz and recapitulated also the known behavior of the variance of the extension fluctuations upon over- and underwinding [59] (Supplementary Figure S5c,d).

### Estimation of labeling efficiency from magnetic tweezers rupture time traces

Examining the extension time traces prior to tether rupture carefully, we find occasional steps in the tether extension (Figure 3a). Increases in tether extension prior to tether rupture are consistent with the disruption of biotin-streptavidin bonds, where the final rupture corresponds to the last biotin-streptavidin pair. We quantify the observed steps and find a rather broad distribution with a mean step size of 13.0 nm ± 14.8 nm (mean ± standard deviation). We then compared the experimental step size distributions (Figure 3b, grey bars) to a simple model for step sizes based on the experimentally used megaprimer sequences and simulated label incorporation (see the section “Model for the step size distributions” in Materials and Methods). Our simple model can account for the overall shape of the observed step size distribution, with an initial sharp increase and subsequent slow decay of the probability of steps with their size. The predicted step size distribution shifts to smaller step sizes for higher label efficiencies *P*_label_ (Figure 3b, colored lines), where *P*_label_ represents a combined effective probability of incorporating a label and binding to an available streptavidin. We find good agreement both of the mean step size and overall distribution (Figure 3c,d) for *P*_label_ = 0.1. The best fitting value of *P*_label_ = 0.1 is 5-fold lower than the ratio of labeled nucleotides in the PCR mix (which is 50%), which suggests that the KOD polymerase used in our experiments preferentially incorporates unlabeled nucleotides, similar to what has been reported for other polymerases [50, 60-64] and/or that not all incorporated biotin labels attach to a streptavidin binding site, likely due to steric constraints. These observations are also consistent with the finding that, while in principle torsionally constrained attachment of DNA or RNA tethers only requires two attachment sites at each end, in practice at least ∼10 potential attachment sites should be incorporated at the respective ends to ensure a high number of stable and torsionally constrained tethers [55].

**Figure 3.**
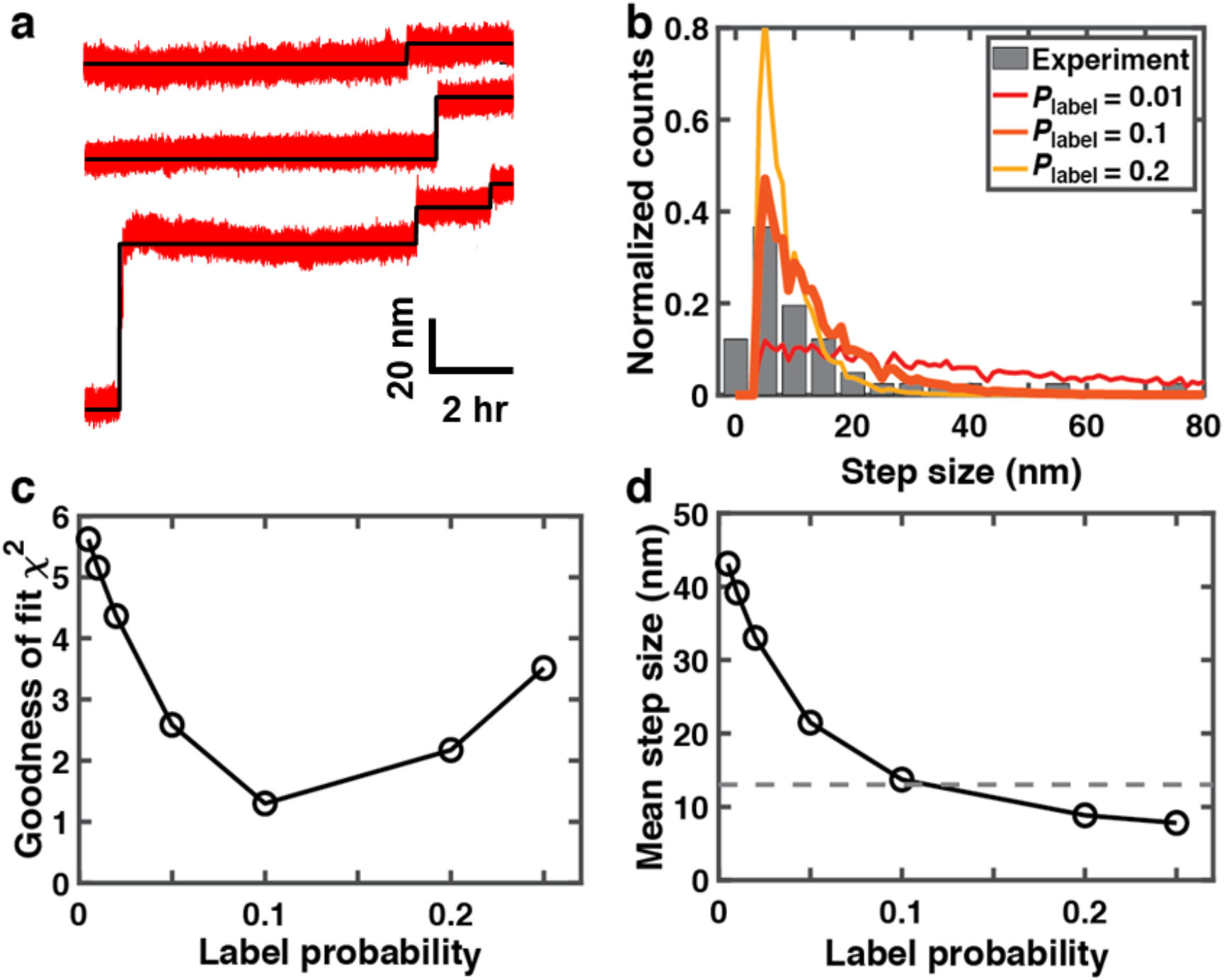
Analysis of tether rupture steps to estimate the labeling efficiency for DNA tethers. (a) Examples of the rupture steps in extension time traces under an applied force of 45 pN (red lines). The black lines are steps in the traces as determined by a step finding algorithm [77]. (b) Experimentally observed step sizes in extension time traces before tether dissociation (gray bars; *N* = 41 steps). Co-plotted are the step size distribution predicted by our simple model (see “Model for the step size distributions” in Materials and Methods) that takes into account stochastic label incorporation for three selected values of the label probability *P*_label_ (color lines; *P*_label_ values are indicated in the legend; simulation results are for 100,000 simulated tethers for each condition). (c) Goodness of fit comparing simulated step size distribution with the experimental data. The goodness of fit was computed as the sum of squared differences of normalized counts in each bin for simulated and experimentally observed data. (d) Mean step size from simulated step size distributions vs. *P*_label_. The horizontal dashed line corresponds to the experimentally determined value. Standard errors of the mean are not shown, as they are much smaller than symbols sizes.

### Assembly of nucleosomes for magnetic tweezers experiments

To test whether our megaprimer assembly-based DNA construct can be used to assemble and measure nucleosomes, we carried out nucleosome reconstitution via salt gradient dialysis to obtain polynucleosomes (Materials and Methods). We use AFM imaging to confirm the assembly of nucleosomes and quantify the different polynucleosome populations, by counting the number of mono-, di-, and tri-nucleosomes that are successfully assembled (Figure 4a). The populations for bare DNA, and DNA with one, two, and three nucleosomes are consistent, within experimental errors, with a simple binomial distribution (Figure 4b), which implies that nucleosome assembly on the three Widom 601 sites is relatively uncooperative under the conditions of our experiments, consistent with previous observations [65].

**Figure 4.**
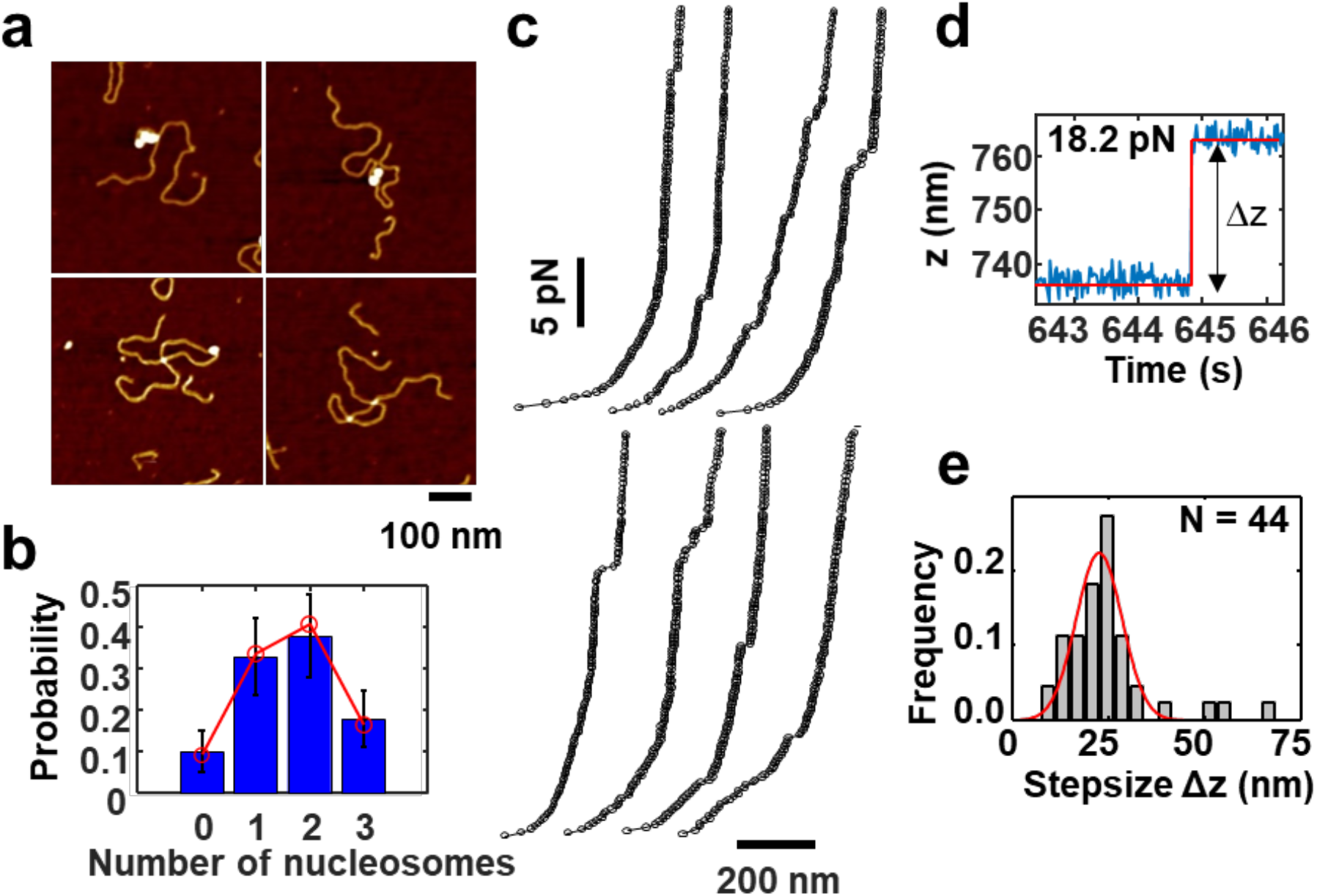
Nucleosome constructs for magnetic tweezers measurements. (a) AFM images of megaprimer PCR generated DNA after nucleosome reconstitution. Images show DNA constructs with 3, 2, 1, or no nucleosomes assembled (left to right, top to bottom). (b) Histogram of the number of nucleosomes assembled on the DNA constructs obtained from AFM images (*N =* 39 DNA molecules; error bars are from counting statistics). Red points are the best fit of a binomial distribution with fitted assembly probability *p =* 0.55. (c) Force-extension curves of polynucleosome DNA constructs from 0.5 to 30 pN. The traces show different length plateaus at forces ≤ 7 pN that indicate outer turn unwrapping and unstacking of polynucleosomes. At higher forces (> 7 pN) we observe discrete steps in extension that represent the full unwrapping of polynucleosomes. (d) Example of a discrete step in time traces at forces > 7 pN, characteristic of the unwrapping of the inner DNA turn from nucleosomes (raw data in blue; the fitted step is shown in red). (e) Histogram of the steps of the inner turn unwrapping. The red line is Gaussian fit, indicating a step size of 22.6 ± 5.9 nm (mean ± standard deviation).

To measure the nucleosomes in the MT, the nucleosome construct is tethered between the bottom surface of a flow cell and a superparamagnetic bead using the same coupling approach that we use for bare DNA (Figure 2a). We performed force-extension experiments on the nucleosome tethers by systematically increasing the force from 0.5 pN to 30 pN in 0.2 pN increments and recording time traces at each force for 5 s (Figure 4c). The extension traces exhibit small and variable steps in the force region between 2-7 pN, which we attribute to unwrapping of the outer turn of DNA from nucleosomes and the disruption of nucleosome-nucleosome interaction. At forces >7 pN, larger and more uniform steps are observed, in agreement with previous reports [36, 66-76] of unwrapping of the inner turn of nucleosomes (Figure 4c,d). We quantify the high-force steps by employing the step finding algorithm by Kerssemakers *et al*. [77] to identify unwrapping steps in our extension vs. time traces (Figure 3d) and to determine the differences of average extensions before and after the steps to get the stepsizes. The distribution of step sizes shows a clear peak at 22.6 ± 5.9 nm (mean ± standard deviation; Figure 4e), in good agreement with previous reports for step sizes of inner turn nucleosome unwrapping in the range of 20-30 nm [36, 66-76].

## CONCLUSION

In summary, we present a new method for preparing functionalized DNA constructs for single-molecule measurements that combines the benefits of ligation-free, high-yield megaprimer-based assembly and covalent attachment of DNA to the surface for very high force stability. Our tethers are torsionally constrained and thus enable torque and twist measurements. In addition, they provide exceptionally high force stability, and we find an average lifetime of (60 ± 3) h at 45 pN using regular, commercially available streptavidin-coated beads. The high yield of correctly labeled DNA enables efficient nucleosome reconstitution by standard salt gradient dialysis, and we demonstrate proof-of-concept measurements on nucleosome assembly quantified by AFM imaging and disassembly under external forces in magnetic tweezers. We anticipate that our methodology will enable a range of measurements on DNA and nucleoprotein complexes that benefit from high yield and force stability.

## ACKNOWLEDGEMENTS

We thank Thomas Nicolaus for laboratory assistance and Lori van de Cauter, Steven De Feyter, Sebastian Konrad, Philipp Korber, and Felix Müller-Planitz for useful discussions.

