## Supplementary Figures S1-S5 for "High-yield, ligation-free assembly of DNA constructs with nucleosome positioning sequence repeats for single molecule manipulation assays"

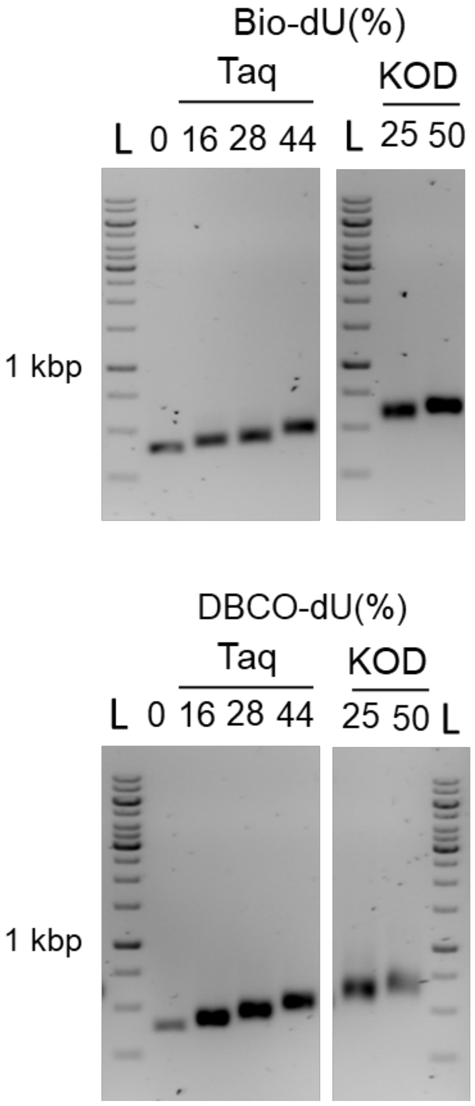


**Supplementary Figure S1. Megaprimers generation by PCR reaction in the presence of labeled nucleotides.** (A) Taq polymerase (NE Biolabs) and KOD Hot Start polymerase (Novagen, Darmstadt, Germany) were used in PCR amplification. Lane L: DNA ladder (1 kb+, NE Biolabs). DBCO-dU, DBCO-$(PEG)_4$-dUTP; Bio-dU, Biotin-16-dUTP; Taq, Taq polymerase; KOD, KOD Hot Start polymerase.

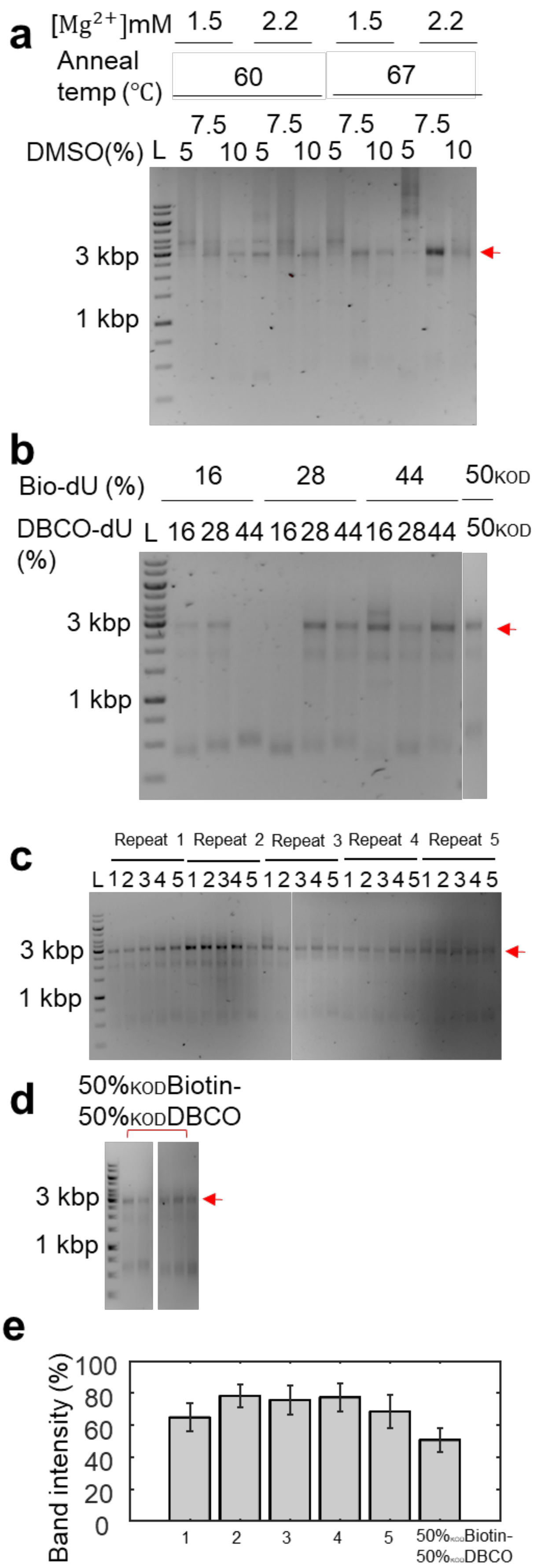
a
[Mg2+]mM 1.5 2.2 1.5 2.2
Anneal temp (℃) 60 67
DMSO(%) L 5 7.5 10 5 7.5 10 5 7.5 10 5 7.5 10
3 kbp
1 kbp
b
Bio-dU (%) 16 28 44 50KOD
DBCO-dU (%) L 16 28 44 16 28 44 16 28 44 50KOD
3 kbp
1 kbp
c
Repeat 1 Repeat 2 Repeat 3 Repeat 4 Repeat 5
L 1 2 3 4 5 1 2 3 4 5 1 2 3 4 5 1 2 3 4 5 1 2 3 4 5
3 kbp
1 kbp
d
50%KODBiotin-
50%KODDBCO
3 kbp
1 kbp
e
Band intensity (%)
100
80
60
40
20
0
1
2
3
4
5
50%KODBiotin-
50%KODDBCO

**Supplementary Figure S2. DNA amplification with megaprimers to produce labeled DNA constructs for single-molecule measurements**. (a) Gel analysis of products of PCR reactions using KOD polymerase to assembly labeled DNA constructs with a linear template with 3 repeats of the Widom 601 sequence and megaprimers generated by Taq polymerase in the previous step. The arrows indicate the expected size of the amplification product. PCR conditions are given in the legend. (b) Products of PCR amplification using KOD polymerase and different ratios of biotin and DBCO labeled megaprimers generated using either Taq and KOD polymerase (rightmost lane). 50KOD; 50% biotin-dU or DBCO-dU labeled prepared using KOD polymerase in the previous step. (c) 5 repeats of PCR experiments using KOD polymerase and Lane 1, 28% Biotin-dU/28% DBCO-dU; 2, 28% Biotin-dU/44% DBCO-dU; 3, 44% Biotin-dU/16% DBCO-dU; 4, 44% Biotin-dU/28% DBCO-dU; 5, 44% Biotin-dU/44% DBCO-dU megaprimers generated using Taq polymerase in the previous step. (d) 5 repeats of PCR experiments using KOD polymerase generated forward and reverse megaprimers. Panels c and d demonstrate the good reproducibility of the megaprimer PCR reactions. (e) Quantification of the target band intensity relative to the integrated total intensity of all bands in the lane of the gels in panels c and d. Numbers refer to the same conditions as in panel c. Values and error bars are the mean and standard deviation from the 5 repeats. We achieve >50% target band intensity for all tested conditions. Image quantification was carried out in Image Lab (BIO-RAD). We selected the relevant lanes and detect band intensities of each lane. The lanes' background were subtracted by specifying the size of a rolling disk (disk size) that determines how closely the background level follows the intensity profile.

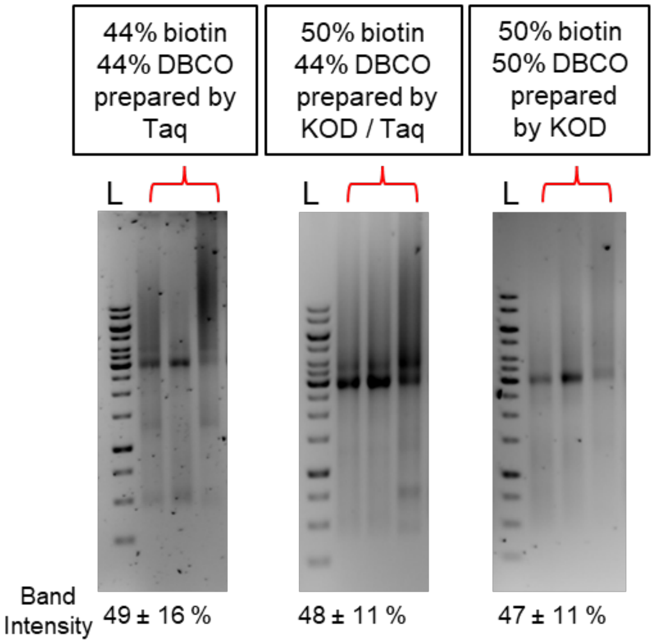


**Supplementary Figure S3. Analysis of megaprimers PCR products obtained in high concentration PCR reactions.** Gel analysis of the products of a modified PCR protocol that uses a smaller final reaction volume and correspondingly higher concentrations of megaprimers and template DNA compared to the protocols described in the main text, to facilitate downstream processing and to test whether this results in a higher yield. Megaprimer conditions are indicated in the panels.

We performed the high concentration/low volume PCR reactions by using 200 ng forward, 200 ng reverse megaprimer, 50 ng linear template, and 10% DMSO in 20 μL 1 x KOD Hot start polymerase Master mix (compared to 100 μl final volume for the reactions described in the main text). We used the following PCR cycling parameters: Initial denaturation at 95 ºC for 2 min; 35 cycles of denaturation at 95 ºC for 20 s, annealing at 60 ºC for 10 s, and elongation at 70 ºC for 65 s. The final cycle was followed by extension at 72ºC for 1 min. We carried out 3 repeats of each PCR reaction with a given set of megaprimer conditions and quantified the relative band intensity of the main product band compared to the integral of all band intensities in one lane. The results suggest that the high concentration/low volume reactions achieve a good yield of the correctly labeled product (visible as the main bands in the gels), but also tend to produce a significant amount of longer products (visible as a smear above the main band). Previous work has suggested that megaprimer concentrations higher than 0.01 μM inhibit the PCR reaction [1]. In contrast, others have argued that megaprimer concentration between 0.02 and 0.04 μM could be advantageous compared to 0.01 μM [2, 3]. We use 0.008 μM megaprimers in the protocol described in the main text and 0.04 μM megaprimer for high concentration / low volume reactions. Our results suggest that both protocols are all suitable for amplifying the final products, but that the larger volume reactions typically have better purity compared to high concentration/small volume reactions, which might be due to the increased tendency for megaprimer mispriming and nonspecific amplification at high concentrations [2, 3]. One advantage of the higher concentration/lower volume reaction is that the subsequent PCR clean up step requires less material and fewer centrifugation steps due to the lower volume.

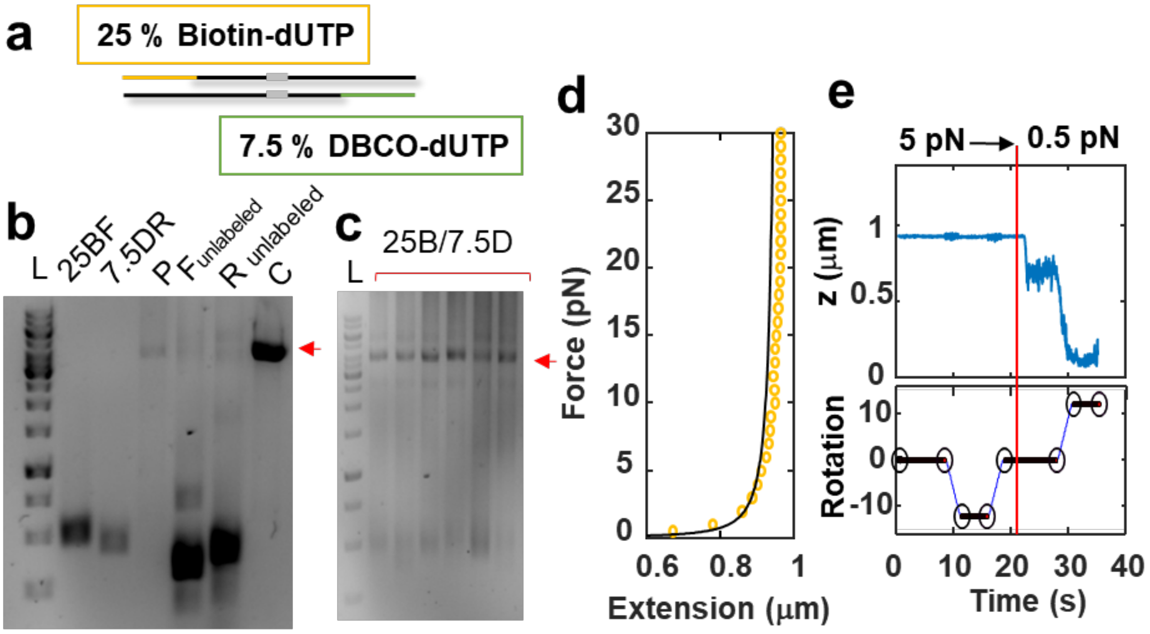


**Supplementary Figure S4. DNA construct with 1x Widom 601 generated with megaprimer PCR reactions.** (a) Schematic of the reaction. The overall approach is the same as in the main text (Figure 1a), but the DNA construct has only one Widom 601 sequence. ~300 bp of both forwards and reverse megaprimer are labeled with, in this case, 25% Biotin-16-dUTP and 7.5% DBCO-(PEG)4-dUTP. We performed PCR reaction with 200 ng forward megaprimers, 200 ng reverse megaprimers, and 50 ng linear template in 20 mL 1x Phusion Hot Start polymerase master mix. We used the following PCR cycling parameters: initial denaturation at 98 ºC for 2 min; 35 cycles of denaturation at 98 ºC for 30 s, annealing at 67 ºC for 30 s, and elongation at 72 ºC for 90 s. The final cycle was followed by extension at 72ºC for 10 min. PCR products were purified by using the QIAquick PCR Purification Kit (Qiagen, Hilden, Germany) after each step of PCR amplification. (b) Gel analysis of the resulting megaprimers and final products. 25BF, 25% Biotin-16-dUTP labeled forward megaprimer; 7.5DR, DBCO-$(PEG)_4$-dUTP labeled reverse megaprimer; P, the final 1 x Widom 601 DNA products made from labeled megaprimer; $F_{unlabeled}$, forward megaprimer without labels; $R_{unlabeled}$, reverse megaprimer without labels; C, the positive control of 1 x Widom 601 DNA without labeled nucleotides. (c) Gel analysis of six repeats of the megaprimer PCR reaction described above. The reaction achieves good yield and reproducibility. (d) Force extension curve of the 1x Widom 601 DNA construct in magnetic tweezers. The black line is a co-plot the inextensible WLC model. (e) Coilability test of the 1x Widom 601 DNA construct in magnetic tweezers. The DNA is twisted first at a force of 5 pN to –12 turns; the fact that the extension does not change indicates correct tethering by a single double-stranded DNA. The DNA is then twisted at 0.5 pN to +12 turns; the fact that the extension decreases indicates that the DNA tether is torsionally constrained.

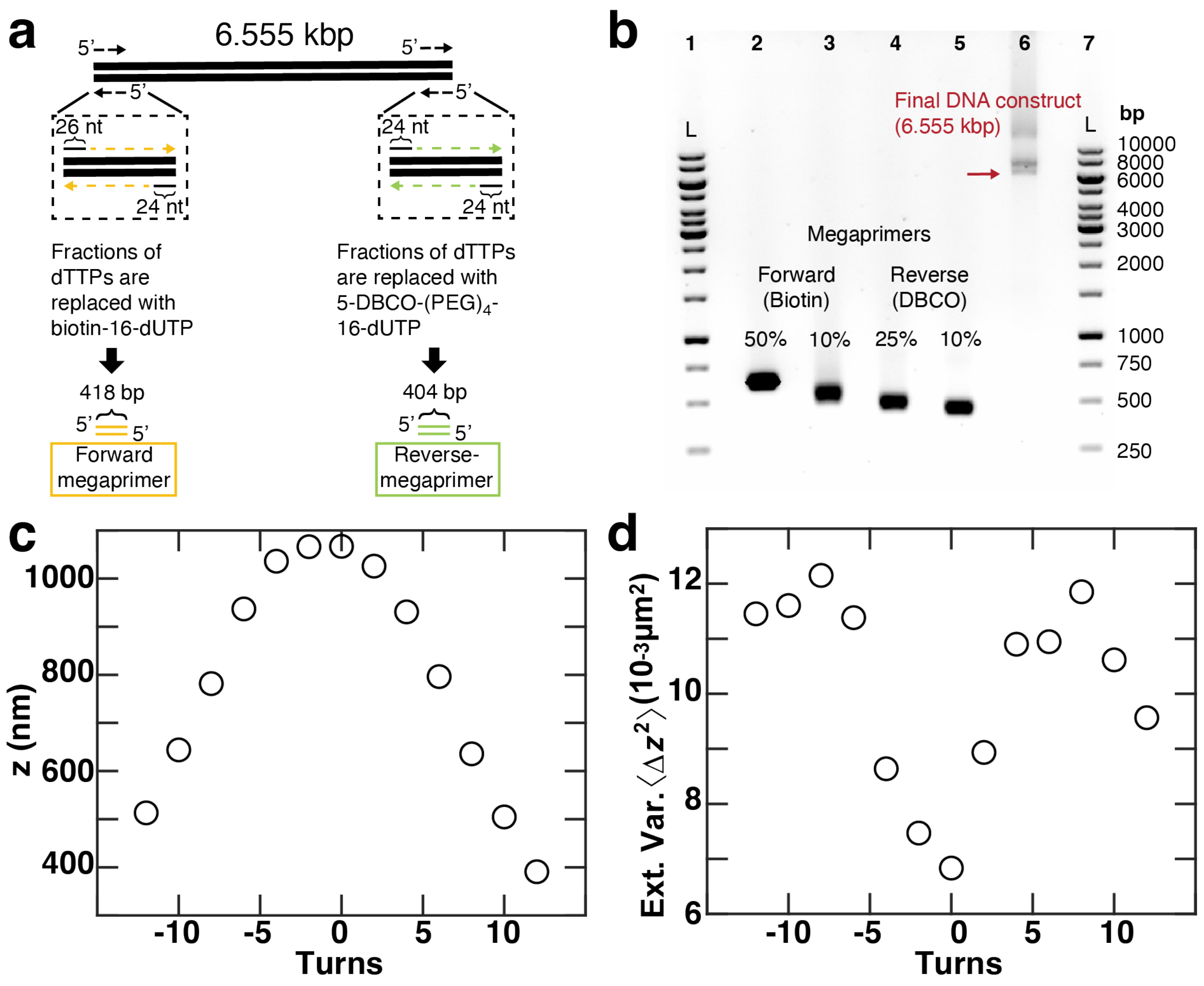


**Supplementary Figure S5. Megaprimers PCR-based DNA assembly of a 6.6 kbp DNA construct and force spectroscopy experiments in magnetic tweezers.** (a) Schematic of the ligation-free megaprimers PCR-based DNA assembly method to synthesize a torsionally constrained 6.6 kbp DNA construct. The overall procedure is the same as what is described in the main text and Figure 1a, however, this reaction uses a DNA without nucleosome positioning sequences and different primer sequences. Two sets of 24-bp (or 26-bp) ssDNA primers and linearized templates are used in PCR reactions to make two multi-labeled 404 bp (or 418 bp) DNAs that become the megaprimers. The two megaprimers are labeled with biotin and DBCO, respectively. A M13mp18-template (NEB; linearized by BspH1 enzyme (NEB)) is used for subsequent PCR to get the final 6555 bp PCR construct. No DMSO was added. (b) Visualization of the PCR result by gel electrophoresis. The left- and right-most lanes (labeled 1 and 7) contain a DNA size ladder (“L”,1 kb gene ruler, NEB). The megaprimers (lanes 2,3,4,5) were generated in PCR reactions using different amounts of biotin-dUTPs and DBCO-dUTPs, respectively, appear at higher bp-values than their actual base pair numbers which is due to the biotin and DBCO labelling that gives them a higher molecular weight. The final DNA construct (lane 6) is indicated by a red arrow. (c) Extension-rotation curve at a constant force of 0.25 pN indicates that the DNA construct is torsionally constrained. Extension data were recorded at 1000 Hz to enable analysis of the variance, in addition to the main extension [4] (d) Variance of the extension of the same bead shown in panel c. The 6.6 kbp DNA construct exhibits the typical response of double-stranded DNA, with decreasing extension and increasing extension fluctuations upon over- and underwinding [4].
